# BET inhibitors synergize with anti-PD1 by rescuing TCF1^+^ progenitor exhausted CD8^+^ T cells in Acute Myeloid Leukemia

**DOI:** 10.1101/2021.08.04.455147

**Authors:** Kyle A. Romine, Hyun-jun Cho, Yoko Kosaka, Kaelan H. Byrd, Jesse L. Coy, Patrick A. Flynn, Matthew T. Newman, Christopher Loo, Jaime Scott, Evan F. Lind

## Abstract

Many acute myeloid leukemia (AML) patients exhibit hallmarks of immune exhaustion, such as increased myeloid derived suppressor cells (MDSCs), suppressive regulatory T cells (Tregs) and dysfunctional T cells. We have developed a mouse model of AML driven by *Flt3-ITD and Tet2* deficiency displays these immune-related features, including CD8^+^ T cells exhibiting a terminally exhausted phenotype (TEx). This T cell subset has been shown to be refractory to immune checkpoint blockade (ICB) monotherapy. Here we show that small molecule inhibitors which target bromodomain and extra-terminal domain (BET) proteins affect both tumor-intrinsic factors but also rescue T cell exhaustion and ICB resistance. *Ex vivo* treatment of cells from AML mice and AML patients with BET inhibitors (BETi) reversed CD8^+^ T cell exhaustion by restoring proliferative capacity and expansion of the more functional precursor exhausted T cells (TPEx). This reversal is enhanced by combined BETi and anti-PD1 treatment. Finally, we show that BETi synergizes with anti-PD1 *in vivo*, resulting in the reduction of circulating leukemia cells, enrichment of CD8^+^ T cells in the bone marrow, and increased expression of *Tcf7, Slamf6*, and *Cxcr5* in CD8^+^ T cells. In total, we show the potential efficacy of combining BETi and ICB therapy in the treatment of AML.

## Introduction

Acute myeloid leukemia (AML) is a genetically heterogeneous myeloid lineage cancer with a 5-year survival of 29% and with limited therapeutic options for those who cannot withstand current frontline therapies^1-3^. The most common mutation, which encompasses 30% of AML patients and is associated with an exceptionally poor prognosis, is an internal tandem duplication in the FLT3 receptor (FLT3-ITD), which leads to ligand-independent signaling to many proliferation pathways. FLT3-ITD is commonly mutated with epigenetic regulators such as those involved in DNA methylation like DNMT3a and TET2^4^. For other difficult-to-treat cancers, such as metastatic melanoma, therapy with immune checkpoint blockade (ICB) has made meaningful increases in life expectancy^5, 6^. ICB functions to reinvigorate cancer-specific T cells via blockade of inhibitory immune checkpoint (IC) receptors which suppress T cell activity and function. Expression of IC receptors, which include CTLA-4, PD1, TIM3, LAG3, TIGIT, VISTA and others, increases after initial antigen exposure and regulates T cell function through various signaling pathways^7^. It has been hypothesized that these receptors are an evolutionary adaptation to chronic antigen exposure to prevent the development of autoimmunity after infection. Recent work has identified unique subsets of CD8^+^ T cells that are generated specifically during chronic antigen exposure. Terminally exhausted T cells (TEx) have been shown to express high levels of IC receptors such as PD1 and TIM3 and low TCF1, while progenitor exhausted T cells (TPEx) are PD1^+^, TIM3^-^ and display high expression of TCF1 and SLAMF6. Importantly, the TPEx population has specifically been shown to expand in response to anti-PD1 therapy and retain anti-tumor capacity^8-12^. In contrast, TEx are significantly more dysfunctional with a lack of proliferative capacity and dramatically reduced secretion of effector molecules, such as granzyme B and perforin^8-10, 13-16^. Our lab and others have recently shown that a proportion of blood and bone marrow specimens from AML patients have an immunosuppressive microenvironment and have hallmarks of immune exhaustion: increased frequencies of regulatory T cells (Tregs)^17^ and myeloid-derived suppressor cells (MDSCs)^18^, decreased T-cell proliferation^19^, elevated expression of immune checkpoint molecules^20-22^ and increased TEx vs. TPEx^22, 23^ populations. Importantly, a subset of these patient samples containing dysfunctional T cells can be rescued by ICB^19^. This prompted us to further investigate potential combination treatment regimens by which immune exhaustion could be reduced in AML.

The Bromodomain and Extra-Terminal domain (BET) protein family is made up of bromodomain-containing proteins BRD2, BRD3, BRD4, and BRDT^24-28^. BET proteins are epigenetic readers which bind acetylated histone residues via conserved BD1 and BD2 domains and mediate downstream functions, such as histone acetylation recognition, chromatin remodeling, and transcription regulation. BRD4 binds acetylated histone tails and recruits Positive Transcription Elongation Factor b (P-TEFb) to enhancer regions to mediate the phosphorylation of the c-terminal domain of RNA polymerase II, required for elongation of the mRNA strand^25, 29^. Loss of function RNAi screening studies identified BRD4 loss as a potent and selective inhibitor of leukemic growth, and treatment with BET inhibitor (BETi) induced leukemia cell death *in vitro* and *in vivo*^24, 27, 28, 30-33^. Clinically, however, BETi therapy was able to elicit some complete remissions but only in a small subset of patients^34^. Interestingly, BETi have also been shown to positively affect CD8^+^ T cells, directly or indirectly, via reduction of PD-L1 expression on myeloid cells^35, 36^ and inhibition of chronic TCR activation genes, such as basic leucine zipper transcription factor, ATF-like (*Batf*), which increased the persistence of stem-cell like memory CD8^+^ T cells^37^. Therefore, we hypothesized that BETi may synergize with anti-PD1 therapy in AML through targeting of tumor-intrinsic factors, such as *myc*, and tumor-extrinsic factors, such as promoting T cell stemness. Accordingly, we showed that a *Flt3-ITD/Tet2* AML mouse model, similar to the previously described *Vav-Cre Flt3-ITD/Tet2* model^38^, phenocopies the immune exhaustion and PD-1 refractoriness found in many patients with AML. We found that these mice have a dramatically increased ratio of TEx:TPEx compared to wild-type (WT) mice and that BETi can rescue immune exhaustion by promoting proliferation of TPEx CD8^+^ T cells. Finally, BETi treatment was found to have a synergistic effect with anti-PD1 *in vivo* and *ex vivo* by reducing leukemic tumor burden and increased RNA expression of TPEx gene programs while decreasing that of TEx. In addition to the mouse model, we observed that BETi rescued T cell exhaustion in a subset of primary AML patient samples.

## Results

### *Flt3-ITD/Tet2* Mice Exhibit hallmarks of immune exhaustion

We previously showed that a genetically-engineered mouse model, *Flt3-ITD*^*+/-*^ *Tet2*^*flox/+*^ *LysM-Cre*^*+/-*^ displays a profound defect in T cell proliferative capacity^39^. To further characterize the tumor microenvironment and functional capacity of T cells in these mice, we assessed changes in myeloid and lymphoid populations at multiple organ sites. We found dramatically increased levels of CD11b^+^ myeloid cells in the spleens and blood of AML mice compared to WT mice (Fig 1a, b). In addition, we observed significant evidence of myeloid infiltration in the spleen causing splenic follicle disruption, and infiltration in liver portal veins and bone marrow (Fig 1c). Furthermore, we found increased levels of GR1^+^ MDSCs and CD4^+^FOXP3^+^ Tregs, a frequent occurrence in AML patients^17-19, 21, 40^(Fig S1a, b), but no difference in T cell frequency (Fig S1c). We also identified significant increases in multiple exhaustion markers in both the CD4^+^ and CD8^+^ T cell populations but most notably a significant decrease in T cell stem-like maintenance transcription factor TCF1 (Fig 1d,e). Taken together, we concluded that these AML mice exhibited a tumor microenvironment characterized by severe immune exhaustion.

**Figure 1:**
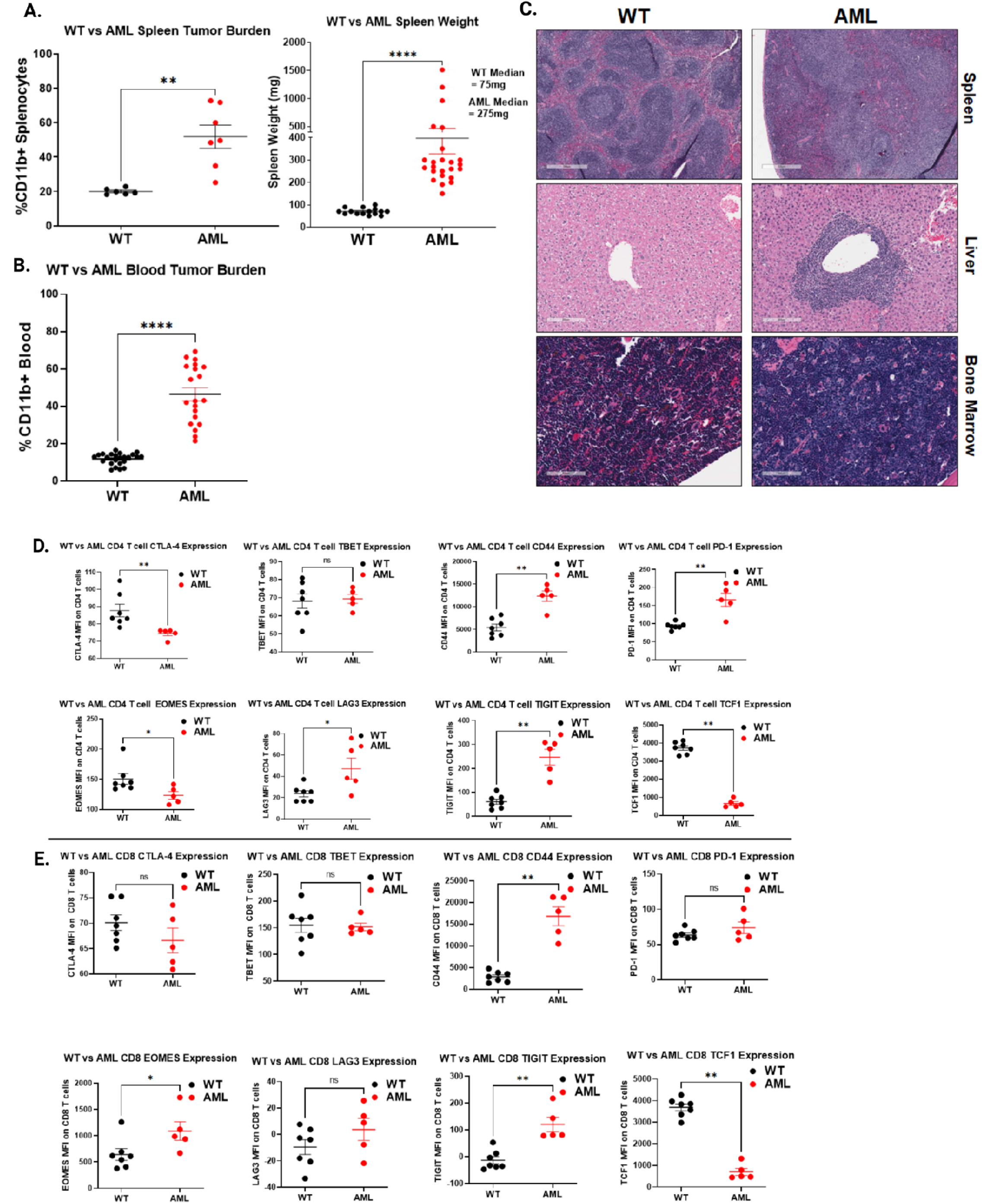
Mouse Model Characterization. a.) Left: Splenocytes isolated from C57BL/6 (WT, black) or *Flt3-ITD +/-, Tet2 +/-, Lys-Cre +/-* (AML, red) mice were stained for live CD11b+ cells and evaluated by flow cytometry. Significance determined by Mann-Whitney T-test. Right: Whole spleens isolated from 15 WT (black) and 24 AML (red) mice were weighed. Median weights for each group are displayed on the right side of plot. Significance determined by Mann-Whitney T-test. b.) Blood isolated from 24 WT (black) and 20 AML (red) mice were stained for live CD11b+ cells and evaluated by flow cytometry. Significance determined by Mann-Whitney T-test. c.) Representative H&E staining of WT and AML mice-derived spleen (Top row, 500 µm scale), liver (middle row, 200 µm scale), and bone marrow (bottom row, 100 µm scale). d.) Splenocytes from untreated WT (black) and AML (red) mice were assessed for expression (median fluorescence intensity) of markers of immune exhaustion on CD4^+^ T cells (Live, CD11b^-^, CD3^+^, CD4^+^). Top row, left to right: CTLA-4, TBET, CD44, PD-1). Bottom Row, left to right: EOMES, LAG3, TIGIT, TCF1). Significance determined by multiple Mann-Whitney T-tests. CD44, PD-1, LAG3 are significantly increased in AML mice. CTLA-4, EOMES, and TCF1 are significantly decreased in AML mice. e.) Splenocytes from untreated WT (black) and AML (red) mice were assessed for expression (median fluorescence intensity) of markers of immune exhaustion on CD8^+^ T cells, as in (d). Significance determined by multiple Mann-Whitney T-tests. CD44, EOMES, and TIGIT are significantly increased in AML mice. TCF1 is significantly decreased in AML mice.

### T cells derived from AML mice are phenotypically and functionally exhausted

We previously showed that T cells from the AML mice are dysfunctional and unresponsive to TCR stimulation ^39^. We assessed the proliferative potential of CD4^+^ and CD8^+^ T cells derived from AML or WT mice by culturing whole splenocytes on anti-CD3 coated plates for 3 days, thus relying on naturally occurring antigen-presenting cells for co-stimulation. As shown previously, both CD4^+^ and CD8^+^ T cells derived from AML mice exhibited significantly reduced proliferation compared to WT mice in response to TCR stimulation (Fig 2a, b). We next asked whether the T cells were intrinsically dysfunctional or if the effect on proliferation occurred only when the T cells were in proximity to tumor cells. We isolated CD3^+^ T cells by negative selection magnetic bead sorting and assessed proliferation with both anti-CD3 and anti-CD28 co-stimulation. The purified T cells from AML mice still lacked proliferative capacity compared to WT, indicating that the T cells are intrinsically dysfunctional (Fig 2c). Finally, we asked whether the AML-derived CD8^+^ T cells were phenotypically terminally exhausted (TEx; PD1^+^, TIM3^+^, TCF1^-^) or TPEx (PD1^+^, TCF1^+^, TIM3^-^), the latter having been shown to retain anti-tumor activity, predict clinical outcome, and expand with ICB^9, 11, 41-44^. Flow cytometric analysis of PD1^+^ CD8^+^ T cells derived from AML mice were primarily consistent with a TEx phenotype, in contrast to WT mice, which were primarily TPEx (Fig 2d,e, S2a). Thus, the AML mice generate an immunosuppressive microenvironment which supports CD8^+^ and CD4^+^ T cell exhaustion.

**Figure 2:**
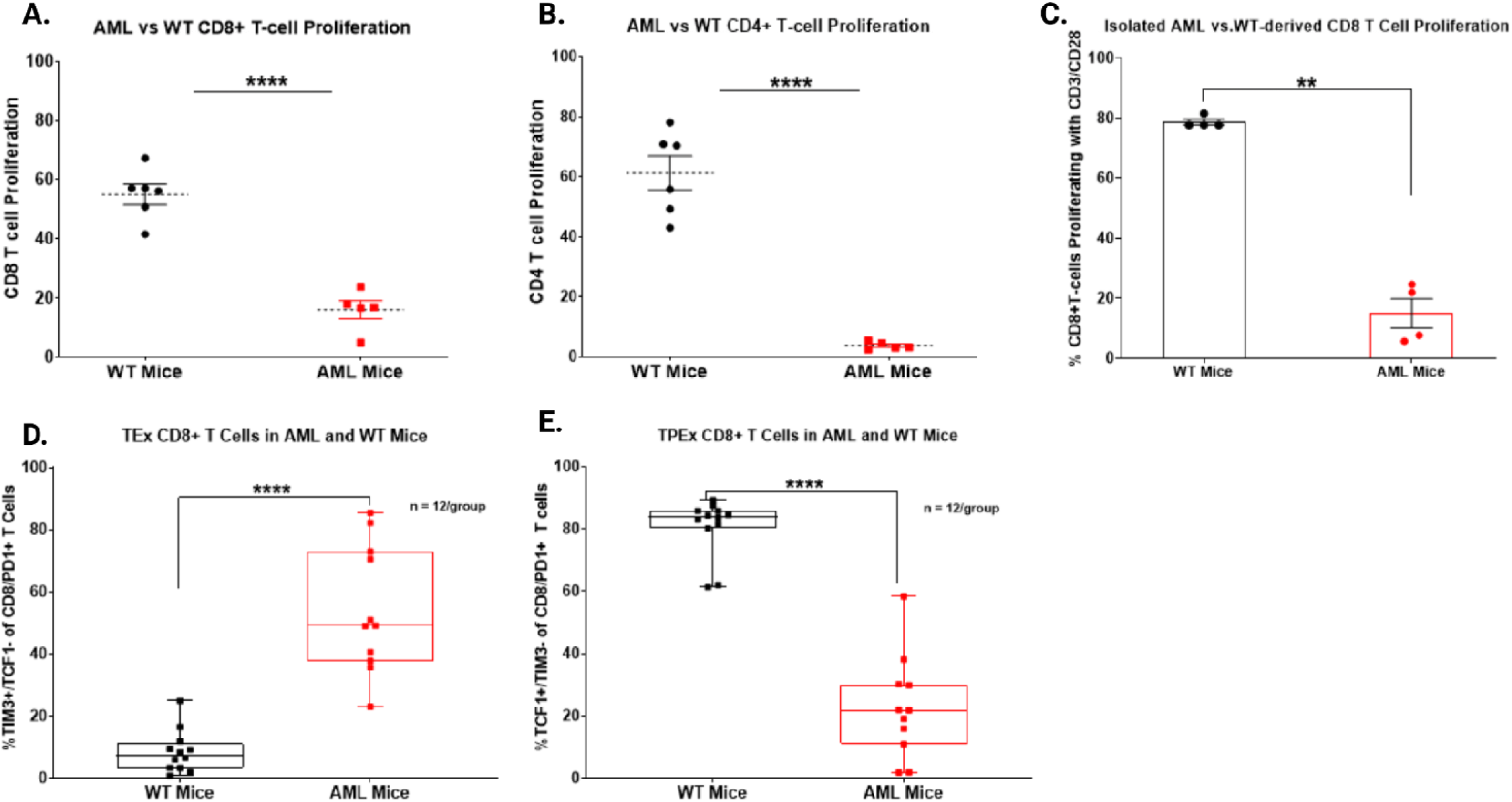
AML mouse-derived CD8^+^ T cells are intrinsically dysfunctional and unresponsive to TCR stimulation. a, b.) WT (black) and AML (red) splenocytes were isolated, stained with proliferation dye (CFSE), and cultured for 72 hours with anti-CD3. Proliferation of a.) CD8^+^ T cells and b.) CD4^+^ T cells was then assessed by flow cytometry, staining with viability and markers to identify T cells. Proliferation displayed is percent CFSE diluted relative to unstimulated (HIgG) control for each cell type. Significance determined by Mann-Whitney T-tests. c.) WT (black) and AML (red) T cells were isolated from splenocytes via CD3 negative isolation magnetic beads. The T cells were then stained with CFSE and plated for 72 hours with anti-CD3 and anti-CD28 stimulation. The cells were then harvested and assessed by flow cytometry. Significance determined by Mann-Whitney T-test. d., e.) Splenocytes derived from WT (black) and AML (red) mice were stained for surface and intracellular markers of T cells exhaustion and evaluated by flow cytometry. d.) Terminally Exhausted CD8 T cells (TEx) are represented as %TIM3^+^/TCF1^-^ of the CD8^+^/PD1^+^ T cells. e.) Progenitor Exhausted CD8 T cells (TPEx) are represented as %TCF1^+^/TIM3^-^ of the CD8^+^/PD1^+^ T cells. Significance determined by Mann-Whitney T-tests.

### *Ex vivo* treatment of splenocytes from AML mice with BETi rescues T cell dysfunction

To identify rationally-derived combination treatment strategies which target tumor-intrinsic and extrinsic factors, we performed high-capacity drug screening as previously described^4^ on tumor cells cultured from the AML mice. Interestingly, we found that three of the top six most efficacious small molecule inhibitors (SMIs) targeted BET proteins (JQ1, OTX-015, CPI-0610) (Fig 3a). Given previous works by Kagoya *et al*., Zhu *et al*., and Hogg *et al*. which established the role for BETi in maintaining stemness of CD8^+^ T cells via inhibition of TCR-activated transcription factor BATF^37^ and that BETi reduce immunosuppressive ligand PD-L1^35, 36^ expression, we asked whether BETi + anti-PD1 could impact the exhausted phenotype found in T cells from our AML mice. To test whether BETi could rescue T cell dysfunction and rescue ICB therapy resistance, we performed proliferation assays as previously described with whole splenocytes and anti-CD3 stimulation but also in the presence of 60nM BETi JQ1, 120nM JQ1, 60nM JQ1 + 10ug/mL anti-PD1, 120nM JQ1 + anti-PD1, or anti-PD1 alone. As expected, treatment with anti-PD1 alone had no effect on proliferation, as the T cells were shown to be primarily terminally exhausted (Fig 2d-g). However, treatment with JQ1 at 60nM, and more so with 120nM, significantly increased proliferation of AML-derived CD8^+^ T cells, whereas WT T cells were unaffected.

**Figure 3:**
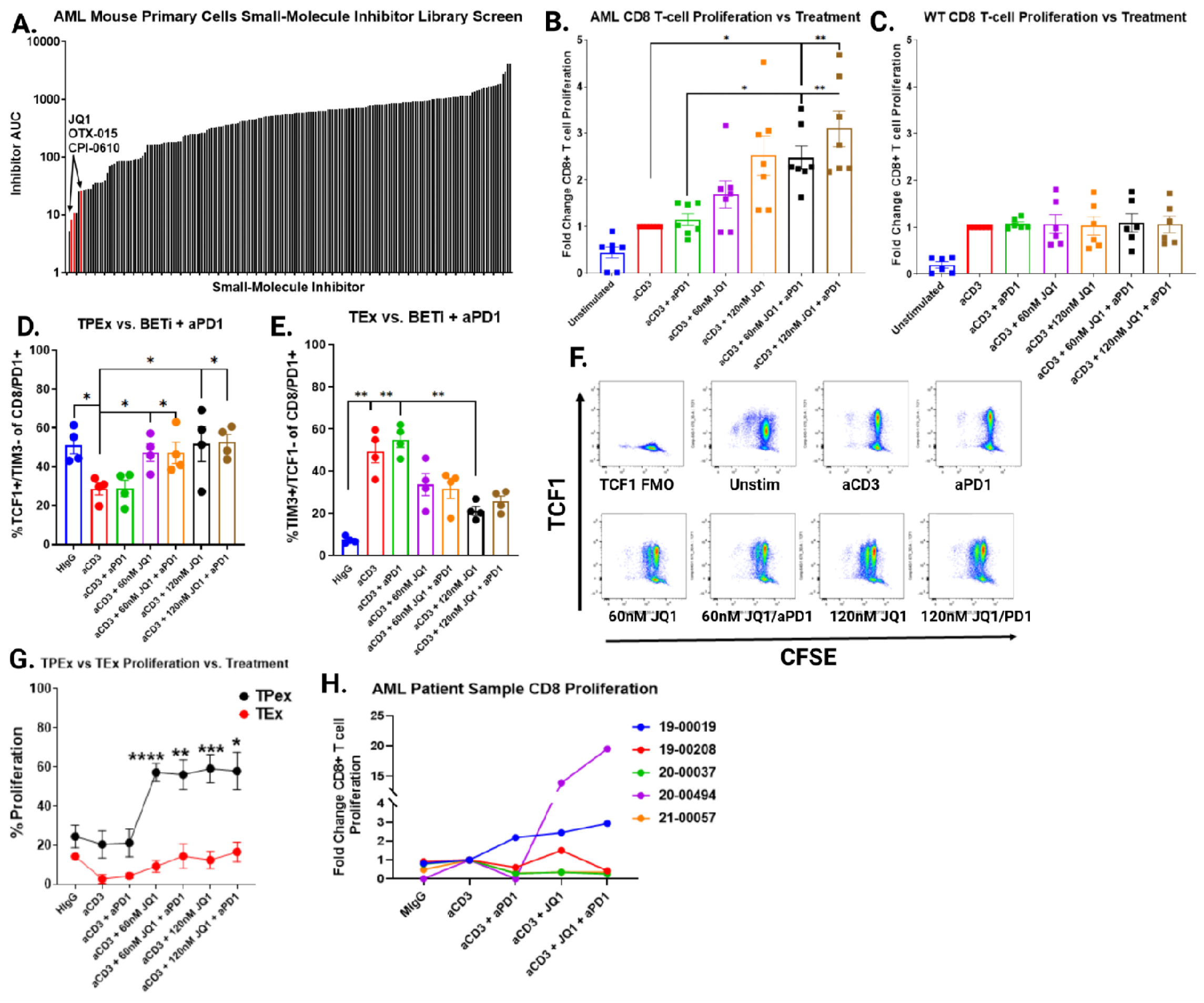
AML mouse T cells are refractory to ICB therapy but partially rescued with BET inhibition. a.) Cells from AML mice passaged for ∼ 1 month were subjected to a small molecule drug library. Cells were seeded into multiple 384 well plates containing titrations of 188 inhibitors, incubated for 72 hours, and viability assessed by MTS assay. Plot represents each Areas Under the Curve (AUC) for every inhibitor on the panel. BET inhibitors JQ1, OTX-015, and CPI-0610 are highlighted in red. b., c.) Splenocytes were isolated from 7 AML (b) or 6 WT mice (c), stained with CFSE, and cultured for 72 hours without TCR stimulation (HIgG), anti-CD3 alone, anti-CD3 with anti-PD1 or titrations of anti-CD3 with JQ1 or anti-CD3 with both JQ1 and anti-PD1. Cells were stained and analyzed by flow cytometry. Plots represent the fold-change in proliferation in CD8^+^ T cells, as measured by percent CFSE diluted relative to anti-CD3 stimulated alone. Significance determined by Kruskal-Wallis multiple comparisons T-tests. d., e.) Effect of BETi, anti-PD1, or BETi + anti-PD1 treatment on the percent of TPEx CD8^+^ T cells (d) and TEx CD8^+^ T cells (e) after culturing for 72 hours as previously in b-c. Significance determined by Kruskal-Wallis multiple comparisons T-tests. f.) Representative dot plots of CD8^+^ T cells from an AML mouse treated as described in 3b,c comparing TCF1 expression vs. CFSE. g.) Splenocytes from 4 AML mice were stained with CFSE and plated for 72 hours without TCR stimulation (HIgG), anti-CD3 alone, anti-CD3 with anti-PD1 and titrations of anti-CD3 with JQ1 or anti-CD3 with both JQ1 and anti-PD1. Proliferation of TPEx (black) and TEx (red) CD8^+^ T cells from AML mice was assessed by flow cytometry. Significance determined by Kruskal-Wallis multiple comparisons t-tests. h.) Fresh mononuclear cells from bone marrow aspirates or peripheral blood obtained from 5 patients with AML were stained with CTV and cultured for 5 days without TCR stimulation (mIgG), anti-CD3, anti-CD3 with anti-PD1, anti-CD3 with 120 nM JQ1 or anti-CD3 with 120 nM JQ1 and anti-PD1. Cells were stained and analyzed by flow cytometry. Plots represent CD8^+^ T cell proliferation for each patient sample.

Moreover, the combination of 120nM JQ1 and anti-PD1 had the largest effect, with a median three-fold increase in CD8^+^ T cell proliferation compared to anti-CD3 stimulation alone (Fig 3b, c). The benefit of adding BETi with anti-PD1 was also observed in CD4^+^ T cells, but was more variable between individual mice, with some mice only marginally benefitting from combined treatment while others were dramatically enhanced (Fig S2b). We next asked what phenotype characterized these newly proliferating CD8^+^ T cells from the AML mice and found that they expressed high levels of TCF1 and low levels of TIM3, indicating that BETi mechanistically act by increasing TPEx:TEx ratios (Fig 3d-g). Moreover, we tested the efficacy of combining BETi + anti-PD1 in five primary AML patient samples with an array of mutational backgrounds. All patient samples were displayed T cell dysfunction. Interestingly, 3/5 samples were at least partially responsive, with one sample dramatically so, to treatment with BETi or BETi + anti-PD1 in both the CD8^+^ and CD4^+^ T cell subsets (Fig 3h, S2c).

### *In vivo*-treated AML mice have reduced tumor burden and increased T cell TPEx gene program expression

Given the results showing the effect of BETi on T cells from the AML mice *in vitro*, we sought to determine the *in vivo* therapeutic efficacy of BETi in combination with anti-PD1 and whether they also modulate the tumor immune microenvironment. AML or WT mice were treated with RIgG (8mg/kg), JQ1 (50mg/kg), anti-PD1 (8mg/kg), or combined JQ1 and anti-PD1 (50mg/kg and 8mg/kg, respectively) for 14 days. We monitored white blood cell (WBC) count in blood pre-treatment, mid-treatment, and at endpoint and characterized the tumor microenvironment (Fig 4a). Measurement of WBC counts over time by hematology analyzer found that only the combination of JQ1 + anti-PD1 significantly reduced tumor burden and, as expected from the *ex vivo* proliferation assays, anti-PD1 alone had no effect (Fig 4b). Characterization of the tumor microenvironment by flow cytometry showed no significant changes to the frequency of Treg cells, MDSCs, or splenic CD3^+^ T cells (Fig S3a,b, 4c). However, we observed a significant increase in CD8^+^ T cells in the bone marrow of mice in the JQ1 + anti-PD1 treatment group, suggesting that the combination has a greater impact on CD8^+^ T cells specifically and not on other immunosuppressive cell types (Fig 4d). Finally, evaluation of RNA-transcripts in isolated CD8^+^ T cells derived from JQ1-treated AML mice identified significantly increased expression of TPEx genes such as *Tcf7, Slamf6*, and *Cxcr5* (Fig 4e). Together, these results highlight the potential of combining BETi and anti-PD1 to treat immunosuppressed T cells in AML, particularly in ICB-refractory cases.

**Figure 4:**
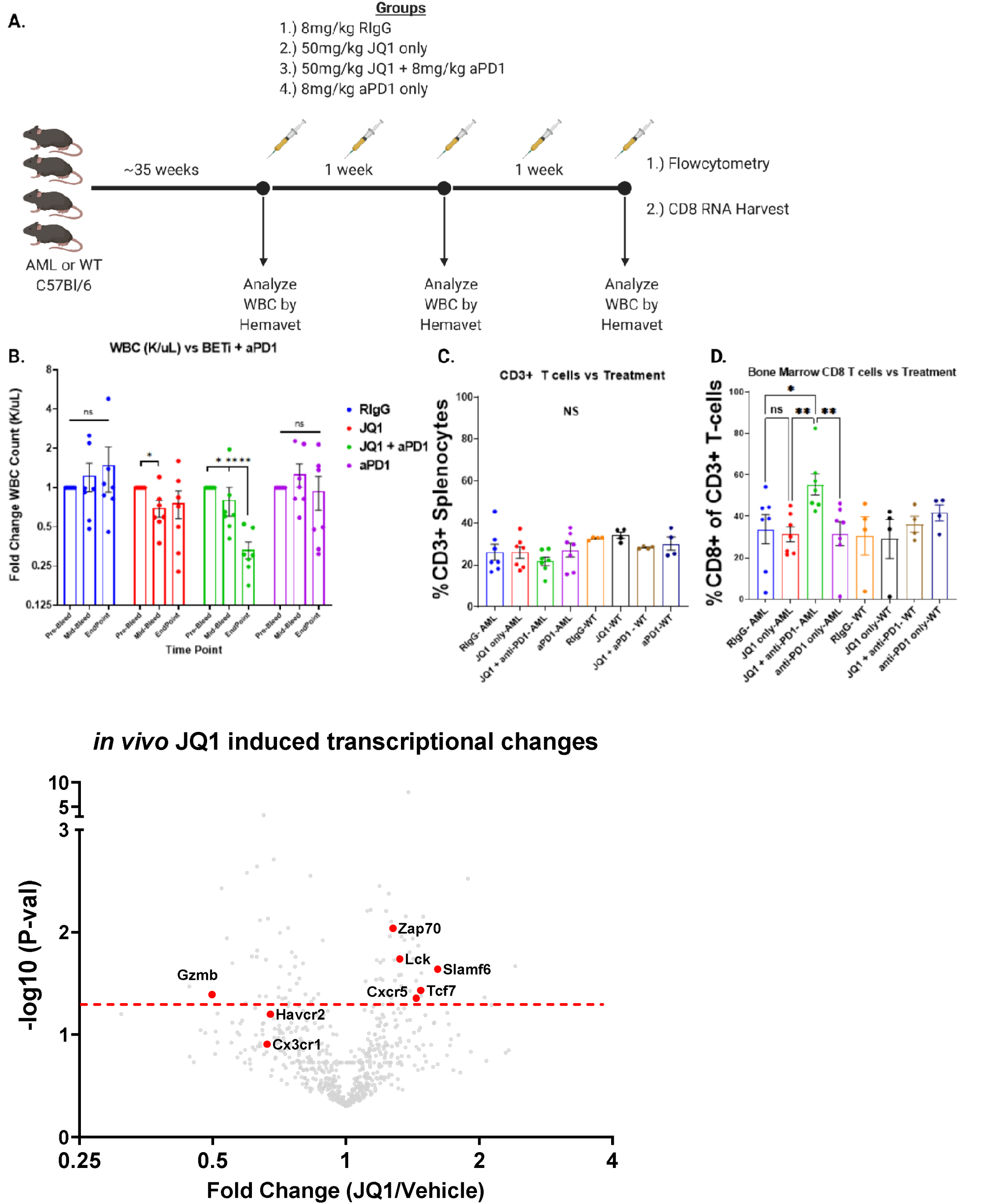
*in vivo* treatment with BETi and anti-PD1 synergizes in reducing tumor burden and enhances CD8+ T cell activity. a.) Schematic detailing *in vivo* BETi + anti-PD1 treatment strategy and functional readouts of efficacy. b.) Mice treated with RIgG, JQ1, JQ1 with anti-PD1, or anti-PD1 alone were bled periodically over a two-week period and assessed for WBC. Data displays fold-change WBC (k/µL) normalized per mouse in comparison to pre-treatment bleed WBC. Significance determined by one-way ANOVA. Timepoints are Pre-Bleed, Mid-bleed (day 7), Endpoint (day 14). c, d.) Bone marrow cells were isolated from treated AML and WT mice and assessed by flow cytometry. Graph denotes %CD3^+^ T cells in the bone marrow (c) and %CD8^+^ T cells in the bone marrow as a percent of all T cells (d). Significance derived from combining two experimental replicates and determined by one-way ANOVA. e.) CD8^+^ T cells were isolated from JQ1-treated and RIgG-treated AML mice and RNA harvested and analyzed by Nanostring. Volcano plot shows the fold-change in normalized transcript levels of JQ1-treated mice vs RIgG treated AML mice vs. –log_10_ P-value as determined by multiple t-tests. Hits of interest are highlighted in red. Dashed red line denotes significance threshold (0.05).

## Discussion

### Immune checkpoint blockade in hematological malignancies

ICB therapy in the blood cancer setting is relatively new and although dozens of clinical trials are ongoing and actively recruiting ^45^, no large-scale study has been completed evaluating the efficacy of anti-PD1 therapies in AML. Our lab previously described the functional immune microenvironment of 50 newly-diagnosed AML patient bone marrow samples^45^ and found that only 41% showed CD8^+^ T cell proliferative capacity that was comparable to healthy donor. Further, we found that the AML patient samples with the most profound proliferative defect (37% of those tested) had severely dysfunctional T cells with significantly decreased production of effector cytokines such as IFN-γ, IL-2, and TNF-α, and increased expression of inhibitory checkpoint markers such as CTLA-4. Of these non-proliferating AML samples, 6 of 18 samples were not rescued with ICB treatment *in vitro*, maintaining the defects in proliferation and effector molecules upregulation seen without ICB. However, 12 of 18 samples did respond, at least partially, to ICB indicating that suppressive mechanisms may be overcome by altering the capacity of T cells to receive inhibitory signals through IC in the tumor environment. Combination treatment regimens that target both the tumor and immune environment to rescue terminally exhausted T cells may be the key to greater success with ICB therapy clinically. Smaller clinical trials have evaluated the efficacy of anti-PD1 + hypomethylating agents (HMA) but only a small subset, approximately 33%, responded to therapy^46^, which is consistent with our *ex vivo* findings^45^. Phenotypic profiling of these responsive patients before and after anti-PD1 +/-HMA identified pre-therapy bone marrow and peripheral blood CD3^+^ and CD8^+^ levels as predictive of response. In addition, we previously described the synergistic combination of combining MEKi and anti-PD1^39^. We found that MEKi acted directly on both tumor cells and immune cells globally by reducing PD-L1 expression on patient samples and, at low doses, restored some proliferative function of AML-derived T cells. These data provide evidence that treatment strategies that target both tumor-intrinsic factors as well as T cell suppression will be more effective.

### Epigenetic regulation of T cell exhaustion

T cell exhaustion is driven by chronic antigen exposure and is accompanied by vast changes in the epigenetic landscape^15, 16, 47-49^. This T cell state is identifiable by three main traits: 1) increased expression of inhibitory receptors such as PD1, LAG3, TIGIT, and CTLA-4, 2) decreased secretion of effector cytokines such as IL-2, IFN-γ, TNF-α and 3) loss of proliferative capacity^16, 50-53^. Epigenetic changes are largely driven by a pool of coordinating T cell receptor (TCR)-responsive transcription factors, such as BATF, IRF4, TOX, and NFAT^8, 51-53^, chromatin remodeling complexes such as EZH2, and other polycomb repressive complex 2 proteins^54-57^.

Generation of TPEx cells relies on the de-repression of many critical pro-memory transcription factors such as TCF1, FOX01, and others as a consequence of targeted DNA methylation deposition acquired during early effector differentiation. Interestingly, the pro-exhaustion transcription factor TOX was found to also directly modulate histone acetylation via direct binding to histone acetyltransferase KAT7^16, 47, 58^. This critical role for epigenetic regulation of T cell exhaustion has garnered much interest in investigating the potential of several epigenetic targeting SMI, whose original design was directed towards targeting tumor-intrinsic epigenetic dysregulation^14^. Very recently, Milner *et al*. showed that BRD4 regulates T cell differentiation by promoting super-enhancer activity at regions regulating key pro-exhaustion/differentiation genes such as *Id2, Cx3cr1*, and *Runx1*^*59*^. Using our unique mouse model of AML, we find that BET inhibition in combination with ICB therapy can lead to a shift from predominantly non-responsive T cells with a TEx phenotype to TPEx and the production of functional CD8^+^ T cells. These results form the basis for further study of such combination therapies in AML. Further studies will focus on understanding the heterogeneity in AML patient responses to BETi + anti-PD1.

## Methods

### AML murine model

Mice expressing FLT3-ITD under the endogenous *Flt3* promotor (strain B6.129-Flt3tm1Dgg/J, The Jackson Laboratory, stock no. 011112) were crossed to mice with the *Tet2* gene flanked by LoxP sites (strain B6;129S-Tet2tm1.1Iaai/J, The Jackson Laboratory, stock no. 017573). The *Flt3-ITD/Tet2* ^*flox*^ mice were then crossed to mice expressing Cre recombinase under the *Lys*m promotor (strain B6.129P2-Lyz2tm1(cre)Ifo/J, The Jackson Laboratory, stock no. 004781). All breeding animals were purchased from The Jackson Laboratory. All mice used in these experiments were bred as heterozygous for FLT3-ITD, TET2, and LysCre. All mouse experiments were performed in accordance with the OHSU Institutional Animal Care and Use Committee protocol IP00000907.

### Flow Cytometry Staining

Bone marrow, blood, or splenocytes were processed and subjected to red blood cell (RBC) lysis by ACK before counting via hemacytometer. Five million cells were resuspended in PBS and stained with Zombie Aqua viability dye (BioLegend, Cat# 423102) for 15 minutes at room temperature, covered from light. The cells were then washed with FACS buffer (PBS, 2% calf serum, 0.02% sodium azide) and resuspended in 25 µL 1:50 mouse FC block (TruStain FcX, BioLegend Cat# 101320), and left on ice for 5 minutes. 25 µL of a 2x cell surface staining antibody cocktail was added directly on top of the cells (final FC block 1:100, 1x Ab concentration) and stained on ice for 30 minutes. For intracellular staining, the cells were then washed with FACS buffer, permeabilized and stained for intracellular targets according to manufacturer’s protocol (eBioscience FOXP3 Transcription Factor Staining Buffer Set, Cat# 00-5523-00), then resuspended in FACS buffer before analyzing on either a BD Fortessa or Cytek Aurora flow cytometer. Data was analyzed using FlowJo software.

### H&E Histology

Liver, spleen, and bone marrow snips were fixed and cut into paraffin blocks for H&E staining. The slides were then scanned using an Aperio AT2 scanner and analyzed with ImageScope.

### Long-term culture of AML mouse-derived bone marrow cells and Inhibitor Library Screen

Bone marrow aspirates from three AML mice were combined, isolated, and cultured in RPMI containing 20% fetal bovine serum (FBS), streptomycin/penicillin, 50 µM 2-mercaptoethanol (RPMI-20), and supplemented with murine SCF (10 ng/ml) and IL-3 (10 ng/ml) (Peprotech or BioLegend). The cells were serially passaged for ∼1 month. Inhibitor library screening to evaluate drug sensitivity was performed as previously described^4^. Briefly, cultured AML mouse-derived cells were counted and seeded into four 384-well plates at a concentration of 2000 cells/well. The cells were then subjected to titrations of 188 unique small-molecule inhibitors in culture for 72 hours. MTS reagent (CellTiter96 AQueous One; Promega) was added and the optical density was read at 490 nm to assess viability.

### Mouse *ex vivo* proliferation assays

96-well round-bottom plates were coated with either 2.5 µg/mL anti-CD3 (Biolegend, clone 145-2C11,Cat# 100359) or HIgG (Biolegend, clone HTK888, Cat#400959), at 4° C overnight, then washed with PBS. Splenocytes were harvested from AML or WT mice and stained with 2 µM CFSE (Life Technologies, Thermo Fisher) at 5×10^6^/mL at 37° C in the dark. After 15 minutes, the CFSE was quenched with 10 mL calf serum and washed with PBS. Cells were then plated at 5×10^5^/well in RPMI with 10% FBS with BME supplementation and penicillin/streptomycin (RPMI-10). After a 72-hour incubation at 37° C the cells were stained for flow cytometric analysis with antibodies listed below and analyzed as previously described. Anti-PD1 (Clone RMP1-14) and RIgG (2A3) were purchased from BioXCell. JQ1 was purchased from Cayman Chemical (#11187).

| Marker | Clone | Vendor |
| --- | --- | --- |
| Tbet | 4B10 | BioLegend |
| LAG-3 | C9B7W | BioLegend |
| TIM3 | RMT3-23 | BioLegend |
| CD44 | IM7 | BioLegend |
| CD62L | MEL-14 | BioLegend |
| PD-L1 | 10F.9G2 | BioLegend |
| CD11b | M1/70 | BioLegend |
| CD8a | 53-6.7 | BioLegend |
| GR1 | RB6-8C5 | BioLegend |
| PD-1 | 29F.1A12 | BioLegend |
| BATF | S39-1060 | BioLegend |
| CD34 | HM34 | BioLegend |
| CD3e | 400A2 | BioLegend |
| TCF1 | C63D9 | Cell Signaling Technologies |
| CD4 | GK1.5 | BioLegend |
| FOXP3 | FJK-16S | BioLegend |

### Human *ex vivo* proliferation assays

Bone marrow aspirates and peripheral blood samples were separated by Ficoll density gradient centrifugation. All experiments were performed using freshly isolated cells. Cells were labeled with CellTrace Violet (CTV, ThermoFisher) and cultured in 96-well plates coated with 1µg/ml anti-CD3 (BioLegend, Clone UCHT1) or control mIgG (BioLegend, Clone MOPC-21). Groups of wells were then treated with either 10µg/ml control mIgG, 10µg/ml anti-PD-1 (EH12.2H7), 60nM JQ1, or 60nM JQ1 + 10µg/ml anti-PD-1. After 5 days, cells were stained for flow cytometry. Viability was determined by Zombie Aqua staining and doublets were gated out of analysis by FSC-A vs FSC-H. Flow cytometry data was acquired on a BD LSRFortessa or Cytek Aurora and analyzed using FlowJo v10 software.

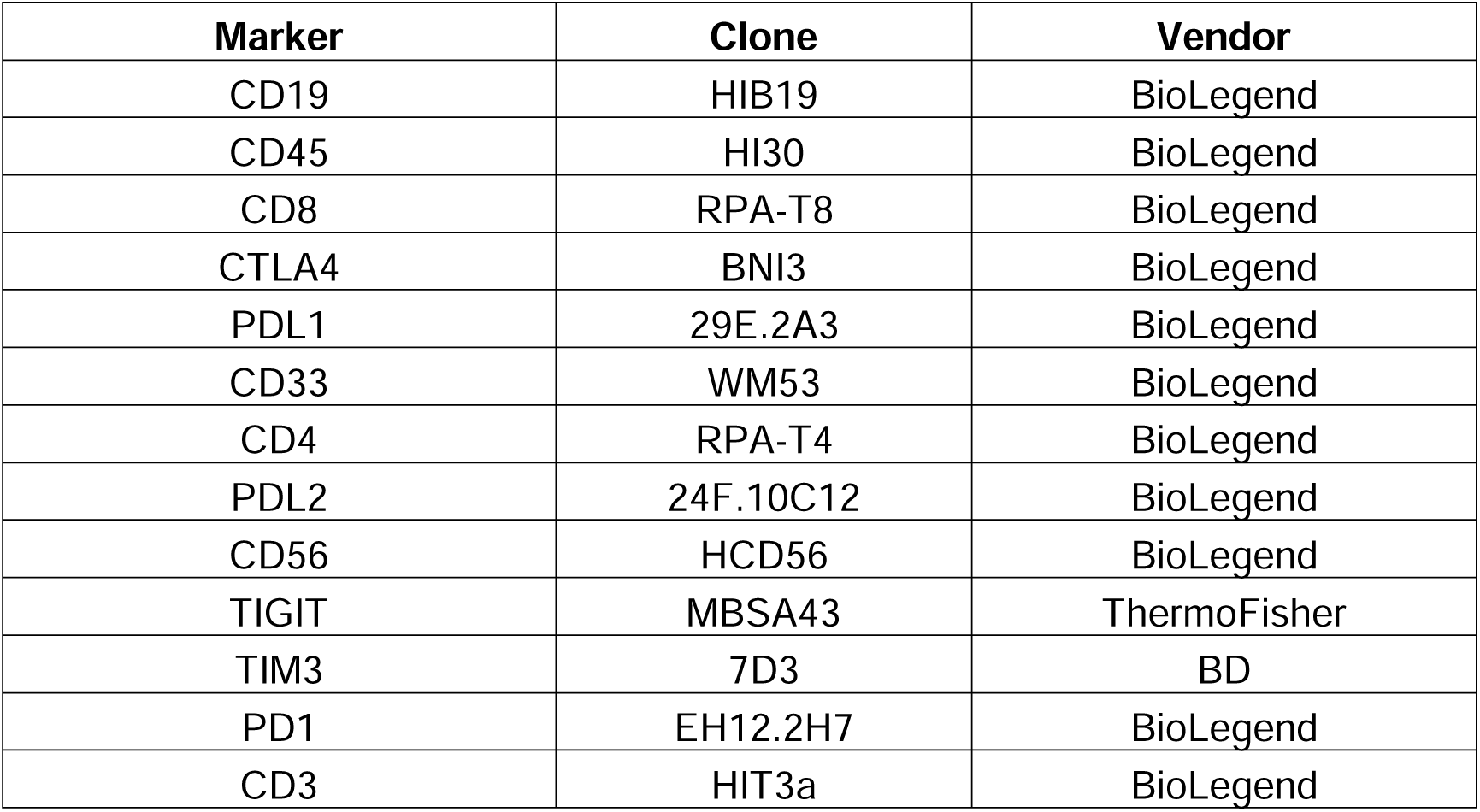

| Marker | Clone | Vendor |
| --- | --- | --- |
| CD19 | HIB19 | BioLegend |
| CD45 | HI30 | BioLegend |
| CD8 | RPA-T8 | BioLegend |
| CTLA4 | BNI3 | BioLegend |
| PDL1 | 29E.2A3 | BioLegend |
| CD33 | WM53 | BioLegend |
| CD4 | RPA-T4 | BioLegend |
| PDL2 | 24F.10C12 | BioLegend |
| CD56 | HCD56 | BioLegend |
| TIGIT | MBSA43 | ThermoFisher |
| TIM3 | 7D3 | BD |
| PD1 | EH12.2H7 | BioLegend |
| CD3 | HIT3a | BioLegend |

### BETi + anti-PD1 treatment *in vivo*

Age range of AML mice used in *in vivo* experiments was 25 to 54 weeks. WT and AML mice were given seven doses of 8 mg/kg RIgG, 50 mg/kg JQ1, 200 8 mg/kg anti-PD1 or 50 mg/kg JQ1 + 8 mg/kg anti-PD1 via i.p injection over two weeks (3 times per week, harvesting 1 day after 7^th^/final injection). Drug solutions were prepared in a solvent of 10% cyclodextrin in PBS from a stock solution of JQ1 (50 mg/mL) in DMSO, followed by sonication/heat bath to dissolve the JQ1. Assessment of tumor burden in the blood, as determined by WBC measurements from Hemavet (950FS, Drew Scientific), was monitored weekly pre-, mid-, and post-treatment. At endpoint, single cell suspensions of bone marrow, blood, and spleen tissues were stained for flow staining as previously described.

### Evaluation of CD8^+^ T cell transcripts via Nanostring

AML mice were given 7 doses of vehicle (10% cyclodextrin) n = 5 or 50 mg/kg JQ1, n = 5, via i.p injection over two weeks (3 times per week, harvesting 1 day after 7^th^/final injection) as previously. CD8^+^ T cells were isolated by magnetic bead isolation (Biolegend #480136) and RNA extracted according to manufacturer’s protocol (PureLink RNA kit, ThermoFisher). Samples were hybridized and loaded onto the Nanostring Chipset according to manufacturer’s instructions (Nanostring Panel NS_MM_CANCERIMM_C3400) and analyzed using the nSOLVER4.0 tool.

## Author Contributions

**K**.**A**.**R**.: Conceptualization, formal analysis, validation, investigation, visualization, methodology, writing-original draft, writing-review and editing. **H-J**.**C**.: methodology, validation **Y**.**K**.: Conceptualization, formal analysis, validation, investigation, writing-review and editing **K**.**B**.: methodology **J**.**L**.**C**.: methodology, writing-review and editing. **P**.**A**.**F**.: methodology, writing-review, editing **M**.**T**.**N**.: methodology **C**.**L**.: methodology **J**.**S**.: methodology **E**.**F**.**L**.: Conceptualization, formal analysis, validation, investigation, writing-review and editing.

## Acknowledgments

This work was funded by a grant NCI U54CA224019 and NCI U01 CA217862 to E.F.L. K.A.R was funded in part by a Program in Enhanced Research Training (PERT) T32 # 5T32GM071338-13. C.L. was funded by NLM 5 T15 LM 7088-28. Flow cytometry was performed at the OHSU Flow Cytometry Core Facility.

## Supplemental Figures

**Supplemental Figure 1:**
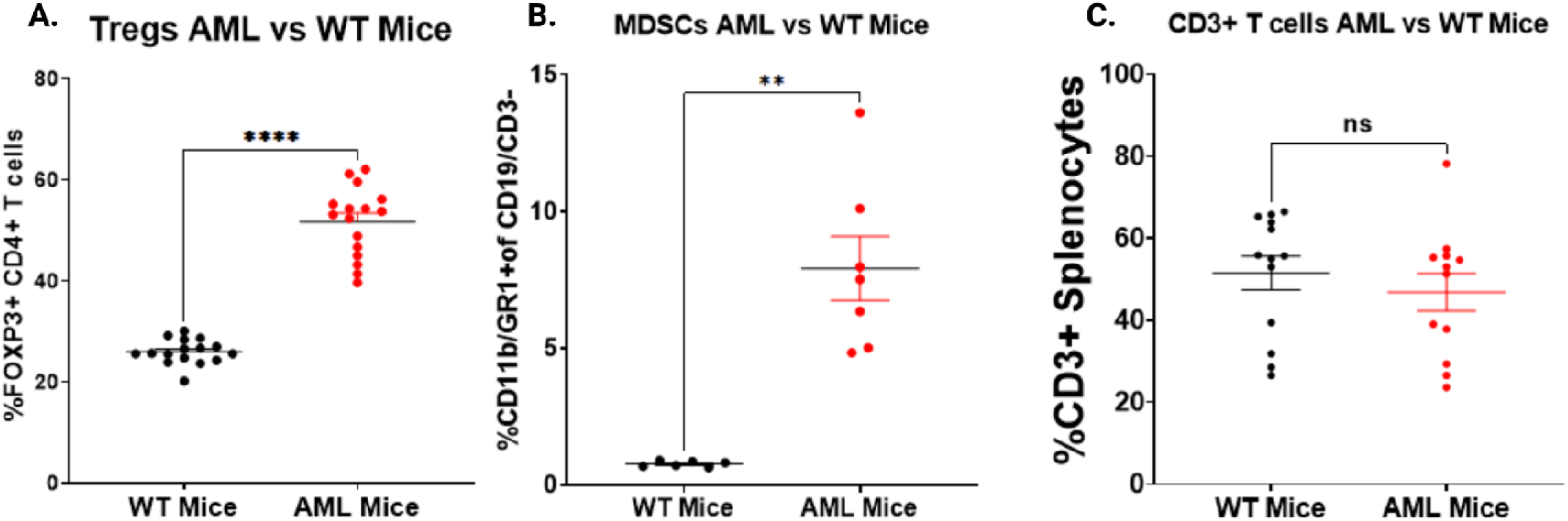
Treg, MDSC, and total T cell frequencies in AML and WT mice. a-c.) Splenocytes isolated from *Flt3-ITD +/-, Tet2 +/-, LysCre +/-* (AML, red) or C57BL/6 (WT, black) mice were isolated and stained for a.) Treg cells (CD4^+^, FOXP3^+^), b.) MDSCs (CD11b^+^, GR1^+^) and c.) all T cells (CD3^+^) and evaluated by flow cytometry. Significances determined by Mann-Whitney t-tests.

**Supplemental Figure 2:**
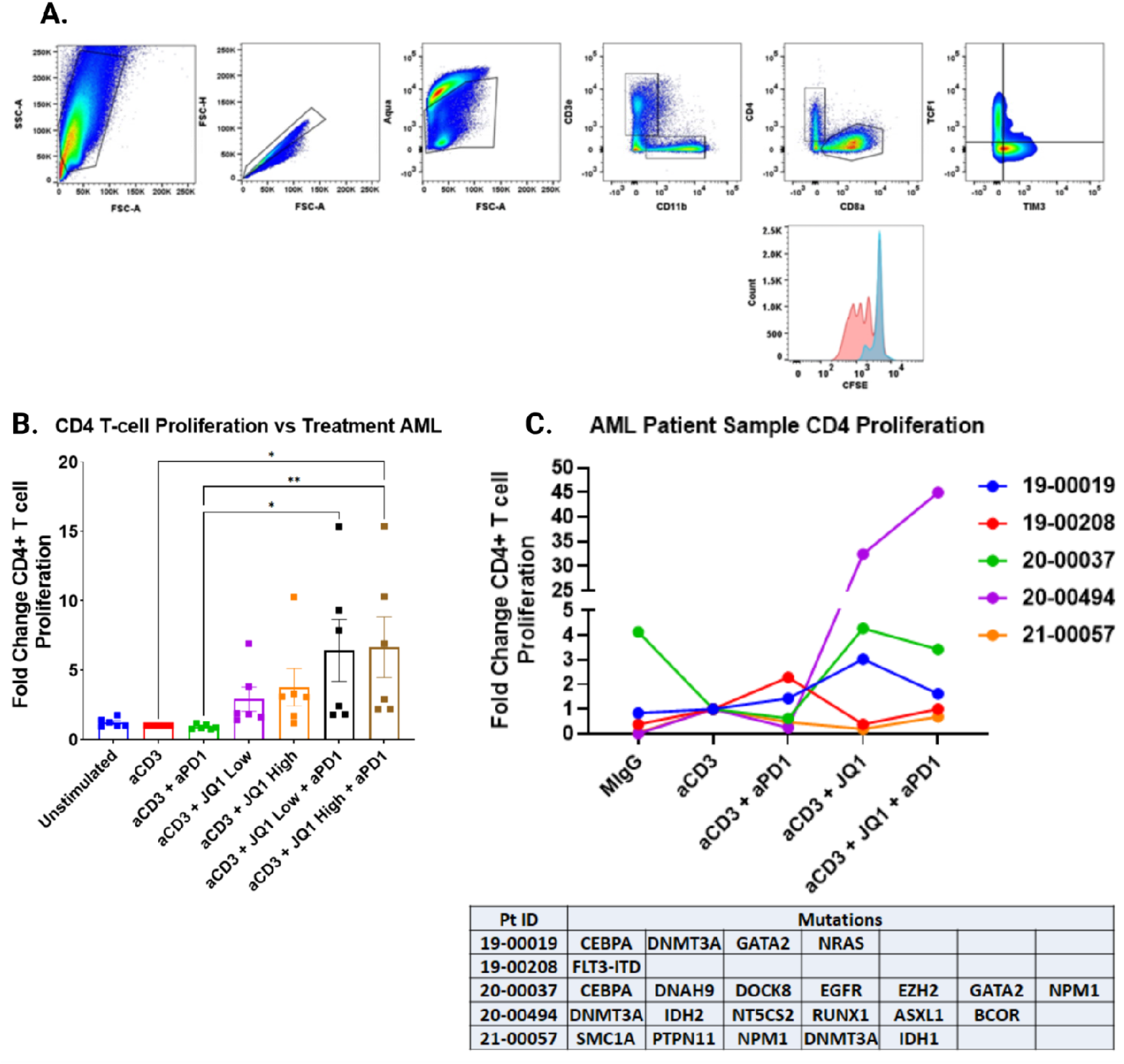
CD4^+^ T cell proliferation with BETi + anti-PD1. a.) Plots below are a representative example from an AML mouse demonstrating the gating scheme used to assess T cell phenotype and proliferation. b.) Splenocytes were isolated from 6 AML mice were cultured for 72 hours without TCR stimulation (HIgG), anti-CD3 alone, anti-CD3 with anti-PD1 and titrations of anti-CD3 with JQ1 or anti-CD3 with both JQ1 and anti-PD1. Cells were stained for assessment by flow cytometry. Plots represent the fold-change in proliferation in CD4^+^ T cells, as measured by percent CFSE diluted relative to anti-CD3 stimulated alone. Significance determined by Kruskal-Wallis multiple comparisons t-tests. c.) Fresh mononuclear cells from bone marrow aspirates or peripheral blood obtained from 5 AML patients were stained with CTV and cultured for 5 days without TCR stimulation (mIgG), anti-CD3, anti-CD3 with anti-PD1, anti-CD3 with 120 nM JQ1, or anti-CD3 with 120 nM JQ1 plus anti-PD1. Cells were then stained for assessment by flow cytometry. Plots represent CD4^+^ T cell proliferation for each patient sample. Corresponding patient sample mutations listed in table below.

**Supplemental Figure 3:**
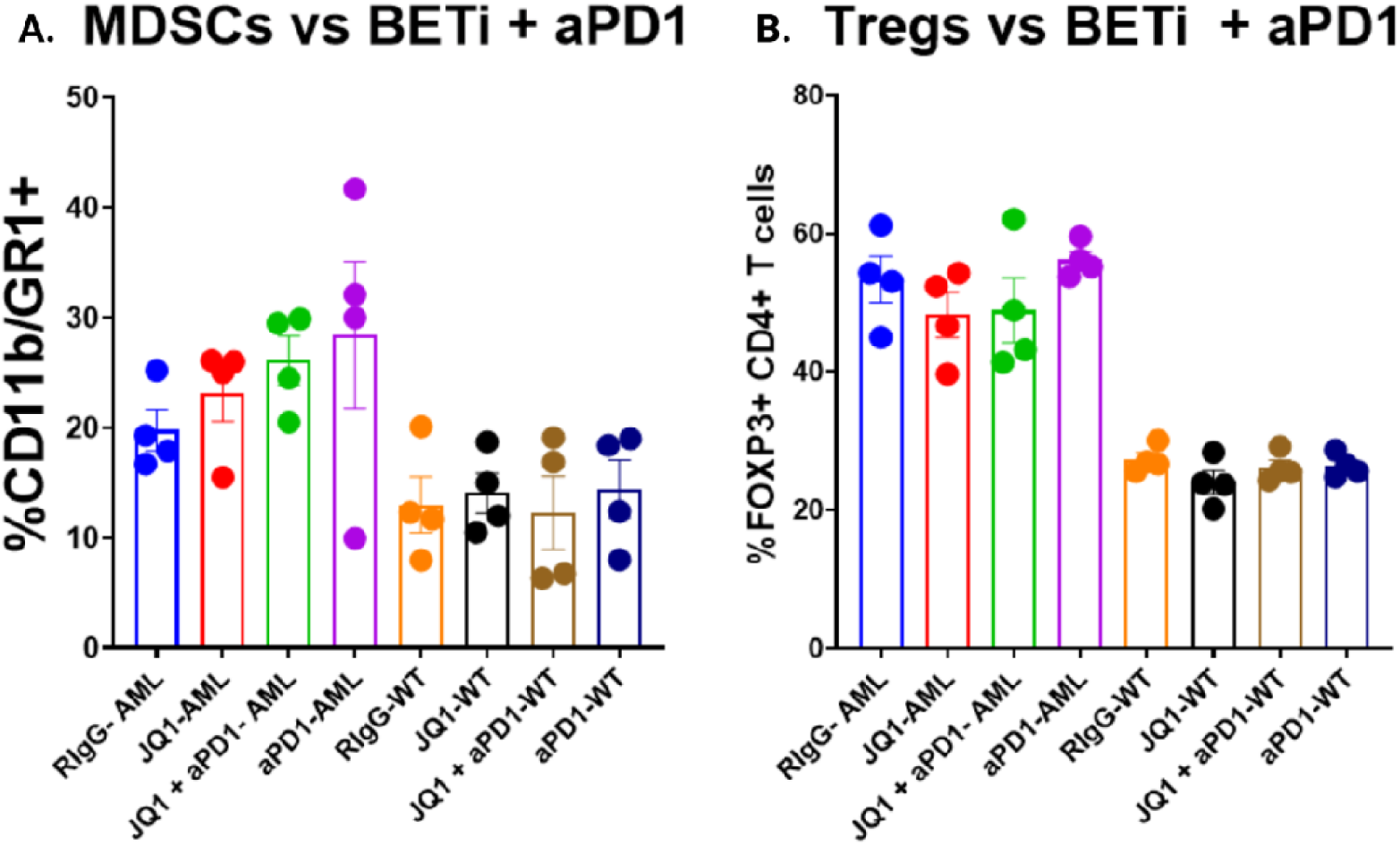
Treg and MDSCs with *in vivo* BETi + anti-PD1 treatment. a., b.) Splenocytes derived from AML mice treated with RIgG, JQ1, JQ1 + aPD1, or anti-PD1 were assessed by flow cytometry to determine a.) %MDSCs (CD11b^+^GR1^+^) and b.) %Tregs (CD4^+^FOXP3^+^). No significance was observed between treatments in AML or WT mice independently.

